# Systematic multi-omics cell line profiling uncovers principles of Ewing sarcoma fusion oncogene-mediated gene regulation

**DOI:** 10.1101/2021.06.08.447518

**Authors:** Martin F. Orth, Didier Surdez, Aruna Marchetto, Sandrine Grossetête, Julia S. Gerke, Sakina Zaidi, Javier Alonso, Ana Sastre, Sylvain Baulande, Martin Sill, Florencia Cidre-Aranaz, Shunya Ohmura, Thomas Kirchner, Stefanie M. Hauck, Eva Reischl, Melissa Gymrek, Stefan M. Pfister, Konstantin Strauch, Olivier Delattre, Thomas G. P. Grünewald

**Author notes:** Correspondence Thomas G. P. Grünewald, MD, PhD, Division of Translational Pediatric Sarcoma Research German Cancer Research Center (DKFZ), Im Neuenheimer Feld 280, 69120 Heidelberg, Germany, Web https://www.dkfz.de/en/translationale-paediatrische-sarkomforschung.

## Abstract

Cell lines have been essential for major discoveries in cancer including Ewing sarcoma (EwS). EwS is a highly aggressive pediatric bone or soft-tissue cancer characterized by oncogenic EWSR1-ETS fusion transcription factors converting polymorphic GGAA-microsatellites (mSats) into neo-enhancers. However, further detailed mechanistic evaluation of gene regulation in EwS have been hindered by the limited number of well-characterized cell line models. Here, we present the Ewing Sarcoma Cell Line Atlas (ESCLA) comprising 18 EwS cell lines with inducible EWSR1-ETS knockdown that were profiled by whole-genome-sequencing, DNA methylation arrays, gene expression and splicing arrays, mass spectrometry-based proteomics, and ChIP-seq for EWSR1-ETS and histone marks. Systematic analysis of these multi-dimensional data illuminated hundreds of new potential EWSR1-ETS target genes, the nature of EWSR1-ETS-preferred GGAA-mSats, and potential indirect modes of EWSR1-ETS-mediated gene regulation. Moreover, we identified putative co-regulatory transcription factors and heterogeneously regulated EWSR1-ETS target genes that may have implications for the clinical heterogeneity of EwS. Collectively, our freely available ESCLA constitutes an extremely rich resource for EwS research and highlights the power of leveraging multidimensional and comprehensive datasets to unravel principles of heterogeneous gene regulation by dominant fusion oncogenes.

## INTRODUCTION

Ewing sarcoma (EwS) is a highly aggressive bone and soft-tissue cancer, mostly affecting children, adolescents, and young adults^1^. EwS tumors are composed of relatively monomorphic undifferentiated small-round-cells^2^. Genetically, EwS is characterized by chromosomal translocations generating in-frame fusions of *EWSR1* and various members of the *ETS* family of transcription factors (in around 85% of cases *FLI1* and in around 10% *ERG*)^3–6^. Different fusion subtypes exist for *EWSR1-FLI1* depending on the number of retained exons of *EWSR1* and/or *FLI1*^3, 5, 6^. The resulting pathognomonic *EWSR1-ETS* fusion oncogenes encode for chimeric transcription factors with neo-morphic features^3, 6, 7^ that rewire the tumor cells’ transcriptome, epigenome, and spliceome (for review see reference^8^). These phenomena are at least in part mediated through EWSR1-ETS binding to GGAA-microsatellites (GGAA-mSats), which are thereby converted into potent neo-enhancers^9, 10^. Apart from *EWSR1-ETS* fusion oncogenes, EwS features a striking paucity of additional somatic mutations and epigenetic alterations in comparison to other malignancies^11–14^.

Despite its monomorphic histology and its rather simple genetic structure, EwS is clinically heterogeneous with diverse outcomes^15–17^. Prior studies based on a limited number of cell line models (all harboring an *EWSR1-FLI1* fusion) suggested that interaction of EWSR1-FLI1 with specific enhancer-like GGAA-mSats and ETS-like binding sites may contribute to EwS tumorigenesis^18, 19^ and clinical heterogeneity of EwS^20^. Indeed, cell lines have been crucial for many basic and translational discoveries in cancer^21, 22^ including EwS^23^. However, the limited number of EwS cell line models assessed to date may have hindered the disclosure of EWSR1-ETS mediated gene regulation principles, and the capture of a broader spectrum of (clinical) heterogeneity, also regarding different types and subtypes of EWSR1-ETS fusion oncogenes. To address this issue, we generated the Ewing Sarcoma Cell Line Atlas (ESCLA), which comprises a panel of 18 molecularly defined EwS cell lines with doxycycline (dox)-inducible short hairpin RNA (shRNA) mediated knockdown of the respective EWSR1-ETS fusion oncoprotein. All 18 EwS cell lines were multi-dimensionally and comprehensively profiled for whole genome, epigenome, transcriptome and spliceome, proteome, and fusion binding to the genome.

## RESULTS

### The Ewing Sarcoma Cell Line Atlas (ESCLA)

The ESCLA (**Fig. 1a**) includes 11 EwS cell lines harboring the most common *EWSR1-FLI1* fusion subtype 1 (fusion of *EWSR1* exon 7 to *FLI1* exon 6)^24^, four with the *EWSR1-FLI1* fusion subtype 2 (fusion of *EWSR1* exon 7 to *FLI1* exon 5), and three being positive for *EWSR1-ERG* (*EWSR1* exon 7 to *ERG* exon 6).

**Figure 1.**
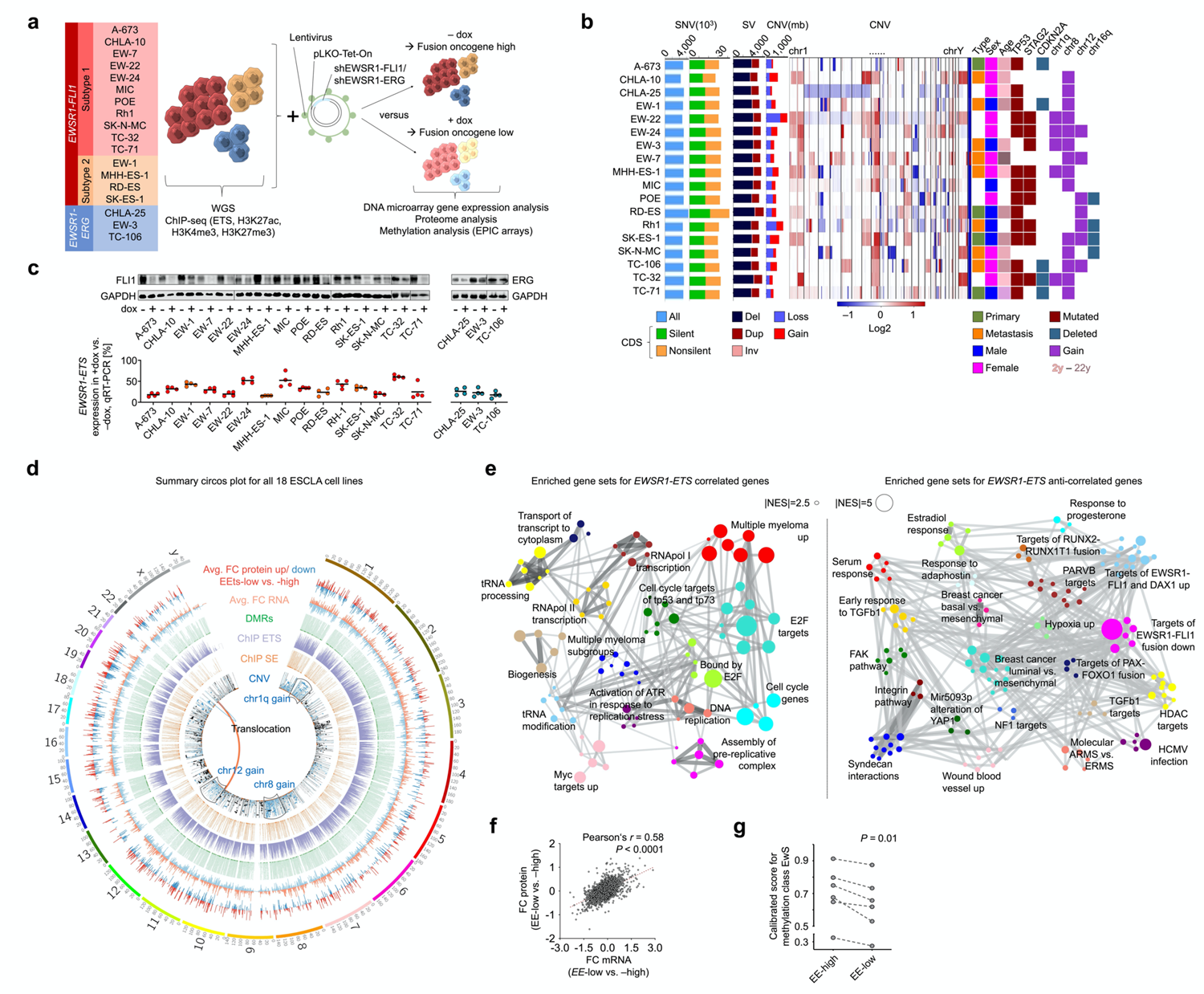
The Ewing Sarcoma Cell Line Atlas (ESCLA) **a)** Schematic of the ESCLA, included cell lines, their fusion (sub)types, and analyses. dox: doxycycline; WGS: whole genome sequencing; ChIP-seq: chromatin immunoprecipitation and subsequent next-generation sequencing. **b)** Bar plots of counts for single nucleotide variants (SNV), structural variants (SV), copy number variants (CNV) per cell line as identified from WGS data, additional heatmap for CNV across the genome and a tile plot indicating clinical characteristics and commonly described variants per cell line. CDS: coding sequence; del: deletion; dup: duplication; inv: inversion; sex refers to the genetical sex of the cell lines determined by copy number analysis of sex chromosomes and STR profiling. **c)** Representative western blot and dot plot for qRT-PCR (*n*=4) for EWSR1-ETS without and with shRNA induction by dox addition, bar indicates mean, GAPDH as housekeeping gene^20^. **d)** Circos plot visualizing genome wide variants, EWSR1-ETS interaction with the genome, and effects of EWSR1-ETS knockdown on transcriptome, proteome and DNA methylation. FC of protein and RNA level are mean values for all cell lines; DMRs, ChIP peak sites and CNVs are displayed stacked for each cell line. Avg.: average; DMR: differentially methylated region; EEts: EWSR1-ETS; FC: fold change; SE: super-enhancer. **e)** Weighted correlation network analysis (WGCNA) on the gene set enrichment analysis (GSEA) results of *EWSR1-ETS*-correlated and -anticorrelated genes at transcriptome level using average rank of expression FC across each cell lines. NES: normalized enrichment score. **f)** Scatter plot indicating average expression FC per gene across cell lines on mRNA and protein level, red line indicates linear regression. **g)** Before-after-plot indicating consistent decrease of the calibrated score for the methylation class EwS (mean value for three replicates, each) in the six cell lines of EwS with highest scores; significance assessed with paired *t*-test.

Whole-genome sequencing (WGS) was performed on all parental cell lines on an Illumina platform with a relatively constant coverage of ∼38x (**Suppl. Fig. 1a**). In accordance with a prior study^4^, the *EWSR1-ERG* positive cell lines (100%) showed evidence for complex chromosomal rearrangements (such as chromoplexy) that could have given rise to the respective fusion^25^. Additionally, 5 of 11 (45.5%) *EWSR1-FLI1* subtype 1 positive cell lines exhibited chromoplexy-typical rearrangements, while the remaining *EWSR1-FLI1* positive cell lines showed characteristics of a balanced chromosomal translocation including an inverse *FLI1-EWSR1* fusion as determined by breakpoint analysis of WGS data and PCR amplification (**Suppl. Figs. 1b–d**)^26, 27^. Moreover, WGS data mirrored previously described common copy number variations, such as gains at chr1q, chr8, and chr12 (present in 28, 78, and 39% of cell lines, respectively)^11, 28, 29^, non-silent mutations in *STAG2* and *TP53* (present in 50 and 78% of cell lines, respectively), and deletions of *CDKN2A* and chr16q in these cell lines (present in 28 and 22% of cell lines, respectively)^11^ (**Fig. 1b**). In addition, our WGS approach using 150 bp, paired-end reads and a PCR-free protocol allowed for genotyping of the mainly intergenic (61.7%) GGAA-mSats with the haplotype inference and phasing for short-tandem repeats (HipSTR) algorithm^30^. Prior studies have described efficient EWSR1-FLI1 binding at GGAA-mSats with ∼13–17 consecutive repeats^18, 20, 31^, and four consecutive repeats as the minimum number for EWSR1-FLI1 binding to those sites^10^. Of the 5,742 GGAA-mSats identified across the hg19 reference genome with min. 4 consecutive to max. 17 GGAA repeats, 3,647 (63.5%) were genotyped for both alleles in our WGS data using HipSTR and high-quality filters (see Methods; **Suppl. Data 1**).

Chromatin immunoprecipitation and subsequent next-generation sequencing (ChIP-seq) was performed with specific antibodies against FLI1, ERG, and the histone marks H3K27ac, H3K4me3, and H3K27me3 in all parental EwS cell lines. ChIP-seq identified in total 156,092 EWSR1-ETS binding sites, of which 91,945 were cell line specific. Although the FLI1 and ERG antibodies did neither differ in the number of ChIP peaks nor in the enrichment of an expected EWSR1-ETS binding site in the ChIP-PCR (1.9**–**8.5% versus 1.6**–**3.5% of input CCND1-EF1 DNA immunoprecipitated, respectively), we noted a relatively high variability of detected EWSR1-ETS peaks across our 18 ESCLA cell lines (median: 13,416 peaks; interquartile range: 24,279). However, considering that especially the cell lines with peak numbers below the median may limit the identification of shared EWSR1-ETS binding sites, we avoided further subclassification of these peaks in ‘weak’, ‘intermediate’ and ‘strong’ using arbitrary cut-offs for subsequent analyses. Yet, despite this limitation, and by analyzing all peaks jointly, we identified 1,888 EWSR1-ETS binding sites that were shared by >80% of EwS cell lines (at least 15 of 18) that were hereafter designated as ‘core subset’ (**Suppl. Data 2**). This core subset included 956 EWSR1-FLI1 binding sites previously identified in two EwS cell lines (A-673, SK-N-MC)^19^, for example the binding sites close to genes being critical for the EWSR1-ETS-mediated transformation capacity, such as *EGR2*, *NKX2-2*, *NPY1R*, *PPP1R1A*, and *SOX2*^18, 32–36^ (**Suppl. Figs. 1e,f**). Using the ESCLA, we identified 932 additional core EWSR1-ETS binding sites, which were adjacent to genes highly significantly enriched in gene ontology (GO) terms involved in developmental and cellular differentiation programs, and in some instances adjacent to other known drivers of EwS malignancy, such as *NR0B1* and *SOX6*^37–39^ (**Suppl. Data 3**).

Genome-wide H3K27ac-based enhancer stitching identified 4,339 super-enhancers (SEs) that were present in at least one cell line. Strikingly, 99.3% (*P*<0.0001) of these SEs exhibited co-localization with EWSR1-ETS binding sites (**Suppl. Data 4**), which supports earlier studies demonstrating a strong effect of EWSR1-FLI1 on the EwS epigenome^19, 40^. Interestingly, 58 EWSR1-ETS-bound SEs that colocalized with 378 genes were shared by all 18 cell lines (**Suppl. Data 4,5**). GO-term enrichment analysis revealed that these 378 SE-associated genes were primarily involved in cellular metabolism and regulation of transcription (**Suppl. Data 5**). These genes also comprised previously reported EWSR1-FLI1 target genes and potential diagnostic markers, such as *PRKCB*, *CCND1*, and *ATP1A1*^41–44^. Even more interestingly, they comprised 17 genes of the developmental *HOX* family of transcription factors (*P*<0.0001) that may play a critical role in Ewing sarcomagenesis^45, 46^.

As expected, peaks of the activating histone mark H3K4me3 rather decorated the transcriptional start site (TSS) of relatively highly expressed genes (median percentile of expression: 70^th^–76^th^ per cell line), while peaks of the repressive histone mark H3K27me3 were found at genes with lower expression levels (median percentile of expression 30^th^–37^th^). Consistently, genes decorated by H3K4me3 were highly significantly enriched for EWSR1-ETS-induced genes and genes decorated by H3K27me3 for EWSR1-ETS-repressed genes (*P*<0.0001, each). 11,090 genes were marked by H3K4me3 in at least 15 cell lines and significantly enriched for GO-terms referring to proliferation, whereas 2,335 genes were marked by H3K27me3 and enriched for GO-terms in the context of neuronal biological processes (**Suppl. Data 6**).

To further investigate the effect of *EWSR1-ETS* on the EwS transcriptome, proteome, and epigenome, all 18 ESCLA cell lines were stably transduced with the dox-inducible pLKO-Tet-On all-in-one system^47^ harboring a fusion transcript specific shRNA for *EWSR1-FLI1* subtype 1 and *EWSR1-ERG* positive cell lines, and FLI1 specific shRNA for *EWSR1-FLI1* subtype 2 cell lines. The knockdown of the respective EWSR1-ETS fusion was validated at the mRNA and protein level by qRT-PCR and western blot, respectively. As shown in **Fig. 1c**, the remaining *EWSR1-ETS* expression upon shRNA induction varied from 15.5**–**60.3% at the mRNA level and was below 30% in 10 cell lines (median: 27.7% across all cell lines).

Interestingly, knockdown of the fusion oncoprotein was accompanied by an increased expression of the wildtype (i.e., non-fused) *FLI1* gene in some *EWSR1-FLI1* positive EwS, even when *FLI1* was targeted by the shRNA. The same effect was observed in 2 out of 3 *EWSR1-ERG* positive cell lines (**Suppl. Fig. 1g**).

Transcriptome analyses were carried out in all cell lines in EWSR1-ETS-high (EE-high) and EWSR1-ETS-low (EE-low) conditions (triplicates per group) using Affymetrix Clariom D arrays (**Suppl. Data 7**) capturing also splice variants. Similarly, proteome profiling was carried out by mass spectrometry, and genome-wide DNA methylation was assessed by Infinium MethylationEPIC BeadChips (Illumina) in EE-high and -low conditions. Collectively, these analyses clearly demonstrated that modulation of *EWSR1-ETS* expression had global and massive effects on the transcriptome, proteome, and methylome across all cell lines (**Fig. 1d; Suppl. Fig. 1h**), which underscores the definition of *EWSR1-ETS* as being ‘dominant’ or ‘master’ oncogenes^20, 32, 48, 49^.

Gene set enrichment analysis (GSEA) and weighted correlation network analysis (WGCNA) revealed that EWSR1-ETS modulated transcripts were most significantly enriched in known EwS-specific and growth promoting gene signatures^8, 48^ (**Fig. 1e**) comprising known EWSR1-FLI1 target genes including *FOXM1*, *LOX*, *MYBL2*^20, 50, 51^. Also, other known EwS fusion targets like *CCK*, *NR0B1*, *PAX7*, *PRKCB*, *SPRY1*, *SOX6* and *STEAP1*^38, 39, 50, 52–55^ showed strong deregulation upon EWSR1-ETS knockdown (**Suppl. Figs. 1i,j; Suppl. Data 8,9**). The *in vivo* differential expression of MYBL2, PAX7, and SOX6 upon EWSR1-FLI1 knockdown was validated at the protein level in tissue microarrays of cell line-derived xenografts comprising 6 of the 18 ESCLA cell lines (**Suppl. Figs. 1k,l**). Notably, mRNA expression regulation by the fusion oncoproteins correlated highly significantly positively with the protein regulation (Pearson’s *r*=0.58, *P*<0.001; **Fig. 1f, Suppl. Data 7,9,10**).

Although one may expect causal epigenetic alterations to precede changes in the transcriptome upon *EWSR1-ETS* knockdown, assessment of the ESCLA cell lines with/without fusion knockdown using Illumina MethylationEPIC arrays did not identify any shared differentially methylated region across all 18 cell lines. In addition, the methylation status of single CpG methylation sites in gene promoters did not significantly correlate with differential gene expression across all 18 ESCLA cell lines (**Suppl. Fig. 1m**). Yet, despite the lack of the natural microenvironment and the 2D nature of the *in vitro* cultured ESCLA cell lines, we noted that they generally clustered in conditions of EE-high with an established reference EwS methylation dataset from 37 primary EwS tumors^56^. The cell lines with the best matching scores were CHLA-10, CHLA-25, MHH-ES1, MIC, RD-ES, and TC-32 (average matching score: 0.685; range: 0.326–0.913). These cell lines showed a significant decrease of the average matching score upon EWSR1-ETS silencing (*P*=0.01; two-sided paired t-test; **Fig. 1g**). A similar observation was made when considering all 18 ESCLA cell lines (*P*=0.002; two-sided paired t-test; **Suppl. Data 11**). These findings may indicate that although EwS tumors exhibit a unique DNA methylation pattern of which part may be caused by the expression of the specific EWSR1-ETS fusion oncoproteins, DNA methylation appears to be not the prevailing epigenetic mechanism through which the transcriptional effects of EWSR1-ETS are mediated^13, 40^. In synopsis, our ESCLA constitutes a high-quality, genome-wide, and multi-dimensional dataset for multiple EwS cell lines representing the major *EWSR1-ETS* fusion types found in EwS (95% of cases).

### Differences and commonalities of distinct EWSR1-ETS fusion subtypes

So far, systemic analyses of potential differences and commonalities of EwS with different types and subtypes of EWSR1-ETS oncoproteins (here EWSR1-FLI1 subtype1, EWSR1-FLI1 subtype 2, EWSR1-ERG) have not been carried out. Hence, we first defined EWSR1-ETS regulated genes and proteins for each cell line separately by a semi-ranked approach in analogy to SE identification with ROSE^57^: To avoid arbitrary cut-offs and false negatives in a *P*-value based approach and to account for variability in the achieved *EWSR1-ETS* knockdown in the given cell line, we ranked all genes by their fold change (FC) of RNA or protein levels (EE-low versus EE-high), respectively, and defined genes/proteins as being differentially regulated when their FC exceeded a cell line specific threshold (see Methods and **Suppl. Fig. 2**). As expected, the calculated thresholds for the RNA level (ranging from –0.478 to –1.585 for downregulated and from 0.468 to 1.614 for upregulated genes upon *EWSR1-ETS* knockdown) correlated significantly with the magnitude of the *EWSR1-ETS* knockdown in the respective cell line (Pearson’s *r*= –0.55 and 0.66, *P*=0.028 and 0.003). Similar observations were made at the protein level (thresholds ranging from –0.357 to –0.915, and from 0.335 to 1.043; Pearson’s *r*= –0.68 and 0.72, *P*=0.002 and <0.001). Applying this method to the multiple cell lines of our ESCLA, profiled on a single platform, enabled a refinement of a previously described EWSR1-ETS-dependent transcriptional ‘core-signature’ that was based (besides various heterologous ectopic *EWSR1-ETS* expression models) on only four different EwS cell lines profiled on different platforms^58^. Defining the refined transcriptional core-signature as concordantly differentially expressed genes (DEGs) in at least 15 of 18 cell lines (>80%) upon EWSR1-ETS knockdown we identified 44 EWSR1-ETS-induced and 26 EWSR1-ETS-repressed core-signature genes (**Suppl. Data 12**). The EWSR1-ETS-induced core-genes comprised *CCK*, *E2F2*, *IL1RAP*, *LOXHD1*, *PPP1R1A*, and *STEAP1*, all of which have been shown to contribute to the malignant phenotype of EwS^36, 53, 55, 59–61^. Conversely, the EWSR1-ETS-suppressed core-genes comprised *LOX* and *FOXO1* that were shown to antagonize EwS malignancy^51, 62^.

In the second step, we compared DEGs across cell lines and fusion subtypes. If the three EWSR1-ETS fusion subtypes represented in the ESCLA would vastly differ in their DNA-binding preferences, the rate of random overlap would be only 3.7% and 4.7% (for mRNA and protein level, respectively)) DEGs across cell lines, even when comparing only the top 33% DEGs of one cell line with all other DEGs of any other cell line. As expected from the very similar molecular architecture of these three EWSR1-ETS fusion oncoproteins and their highly conserved C-terminal DNA-binding domains^6, 63^, we noted a rather large and highly significant overlap in DEGs at both the mRNA and the protein level of 23–41% (*P*<0.001 for both; **Figs. 2a,b**). In support of this finding, GSEA on gene lists ranked by their mRNA or protein regulation by the different fusion oncoproteins, resembled each other for the different fusion oncoproteins (**Figs. 2c,d, Suppl. Data 13**). Furthermore, no subgroups related to the respective fusion oncoprotein could be identified among the 18 ESCLA cell lines when applying t-distributed stochastic neighbor embedding (t-SNE) to the mRNA expression, protein expression, or CpG methylation (**Fig. 2e–g**). Yet, the observed overlaps at the transcriptional and proteomic levels were far from being perfect, even when comparing DEGs between cell lines with the same fusion subtype, suggesting that factors other than the fusion subtype may cause heterogeneity of DEGs.

**Figure 2.**
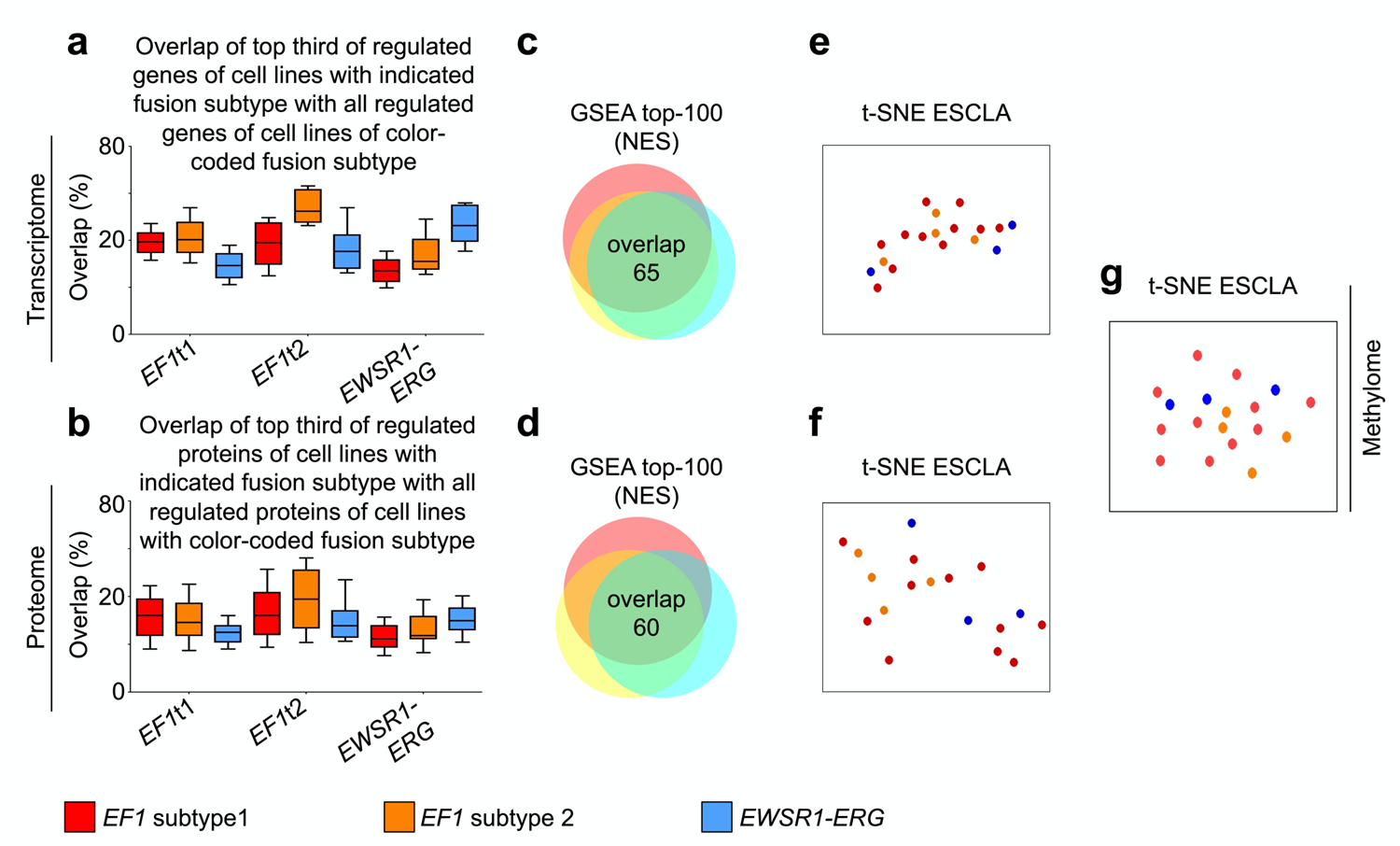
Differences and commonalities of distinct EWSR1-ETS fusion subtypes. **a)** Box plot indicating overlap of the top third genes regulated on mRNA level upon EWSR1-ETS knockdown (only those 924 genes with highest fold changes were considered as regulated, as this was the minimal number of regulated genes among all cell lines; the top-third corresponds to the 208 genes of the 924 with highest fold changes) of any cell line positive for the fusion indicated on the x-axis with the top 924 regulated genes of all other cell lines with the color-coded fusion (sub)type. EF1t1/2: EWSR1-FLI1 subtype 1/2. **b)** Box plot as in **a** for the top-third (72) regulated proteins of any cell line with fusion (sub)type indicated on x-axis versus the top 216 regulated proteins (minimal number of regulated proteins among all cell lines) of all other cell lines. **c)** Venn diagram indicating overlap of the top 100 GSEA results per fusion type; GSEA was performed on gene list ranked by the average fold change rank of each gene across all cell lines of a specific fusion sub(type) on mRNA level and **d)** on protein level. NES: normalized enrichment score. **e)** and **f)** t-distributed stochastic neighbor embedding (t-SNE) plots for the full transcriptome and proteome of all 18 EwS cell lines in the ESCLA. **g)** t-SNE plot for all 18 EwS cell lines in ESCLA based on DNA methylation.

### Characteristics of EWSR1-ETS-bound GGAA-mSats in the ESCLA

While the mechanisms through which EWSR1-ETS fusions suppress the expression of specific genes are poorly understood and likely mediated indirectly, a direct transactivational role is considered for the majority of EWSR1-ETS-induced genes. For induction of gene expression, EWSR1-ETS binding to GGAA-mSats has been shown to play a pivotal role in EwS^9, 18, 34, 38, 39^. Consistently, we found that EWSR1-ETS-induced genes, as identified by the rank-based cell line-specific approach, were strongly and highly significantly enriched for nearby EWSR1-ETS-bound GGAA-mSats (defined as at least four consecutive GGAA repeats) compared to unregulated genes (on average 1.41 versus 0.98 EWSR1-ETS-bound GGAA-mSats in a window of 2 Mbp around the TSS, *P*<0.0001) in the ESCLA. In agreement with a rather indirect regulation of genes suppressed by EWSR1-ETS (i.e., genes being upregulated upon EWSR1-ETS knockdown), such an enrichment of EWSR1-ETS-bound GGAA-mSats was less pronounced for those genes (on average 1.11 bound GGAA-mSats).

Since GGAA-mSats are highly polymorphic^38, 64^, we hypothesized that their polymorphism may affect EWSR1-ETS binding and, thus, result in heterogenous (neo-)enhancer function on EWSR1-ETS regulated genes. To test this hypothesis, we carried out several analyses to assess the ‘length’ and composition of EWSR1-ETS-bound and -unbound GGAA-mSats as well as their flanking regions across our ESCLA. Since earlier reports have shown an essential role of the number of consecutive GGAA repeats^9, 18, 20, 39^, we first studied this aspect based on the HipSTR output. As shown in **Fig. 3a**, both the mean GGAA repeat number of both alleles as well as the number of GGAA repeats of the ‘longer’ allele per locus were strongly and highly significantly correlated with the rate of observed EWSR1-ETS binding in the respective cell line (Pearson’s *r²*=0.98 and 0.99, *P*<0.0001). In agreement with studies on single cell lines describing a ‘sweet spot’ for EWSR1-ETS binding^9, 38^, we noted a second maximum of genotyped GGAA-mSats (for both mean and maximum allele length at around 12–14 consecutive GGAA repeats). Interestingly, the increase of the EWSR1-ETS-binding rate over the count of maximum consecutive GGAA repeats of both alleles had a slight latency when compared with the average count, indicating that long consecutive GGAA-mSats on both alleles increase the probability of EWSR1-ETS-binding compared to loci containing only one ‘long’ allele (min. mean allele length versus max allele length with >50% EWSR1-ETS binding 12 versus 13 GGAA repeats, **Fig. 3a**). Next, we assessed the impact of additional GGAA motifs and ‘shorter’ GGAA-mSats and the nature of interspacing bases around the longest consecutive GGAA stretch on EWSR1-ETS binding. These additional GGAA motifs and bases were also genotyped by HipSTR within the GGAA-mSats’ genomic coordinates predicted by Tandem Repeats Finder^65^, and their rates of occurrence at EWSR1-ETS-bound and -unbound GGAA-mSats were calculated. To reduce noise, only homozygous alleles were investigated, and the calculated ratios were normalized with the expected ratio for GGAA-mSats with the same number of consecutive motif repeats, but without further genotyped bases. As shown in **Figs. 3b,c**, GGAA-mSats that were often bound by EWSR1-ETS exhibited, besides relatively high number of GGAA repeats at the ‘longest’ stretch, typically the following two related characteristics: I) a high number of additional single GGAA motifs and/or additional GGAA repeats in their vicinity, and II) a low number of interspersed bases or flanking bases to the next GGAA motif/repeat.

**Figure 3.**
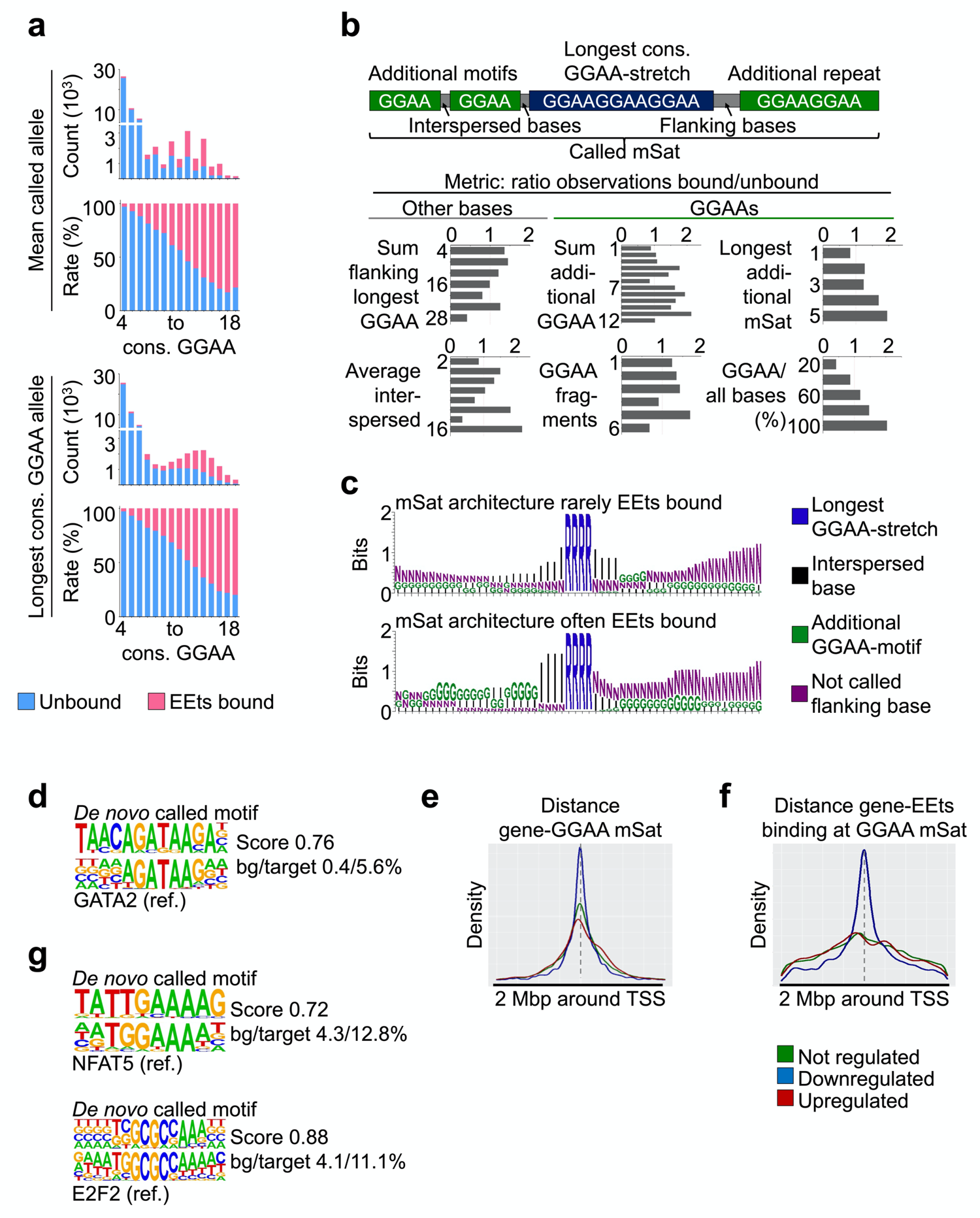
Characteristics of EWSR1-ETS-bound GGAA-mSats in the ESCLA: **a)** Bar plots indicating absolute and average count of genotyped GGAA-microsatellites (mSats) stratified per average (upper panel) or maximum (lower panel) consecutive (cons.) GGAA-repeats for both alleles, EWSR1-ETS (EEts) binding is indicated with pink color. **b)** Top panel: Graphical description of the terms additional motif, longest GGAA-stretch, flanking and interspersed bases and addition repeats in GGAA-microsatellites (mSats) genotyped by HipSTR, here employed to describe the GGAA-mSat architecture. Below: Bar plots indicating the ratio of GGAA-mSats found EWSR1-ETS-bound/-unbound in ChIP, normalised to the expected ratio for the longest GGAA-mSat, stratified by different components of the GGAA-mSat architecture. **c)** Motifs for the most often EWSR1-ETS (EEts)-bound and -unbound GGAA-mSats. **d)** *De novo* called motif in the flanking region of often EEts-bound GGAA-mSats and matched known reference binding motif for GATA2. bg: background (i.e. control sequences) frequency of the *de novo* motif; target: *de novo* motif frequency in investigated target sequences; ref: reference binding motif. **e)** Density histogram for the distance of a GGAA-mSat to the transcription start site (TSS; indicated as interspersed grey line) of a downregulated (blue), upregulated (red) and unregulated (green) gene upon EEts-knockdown; genes were considered as down- or upregulated when the regulation was observed in at least 33% of cells. **f)** Density histogram indicating distance of the next EEts-bound GGAA-mSat to the transcription start site (TSS; represented as interspersed grey line) of a downregulated (blue), upregulated (red) and unregulated (green) gene upon EEts-knockdown in the respective cell line. **g)** *De novo* called motifs for transcription factor binding in the promoters of EEts-regulated genes.

Finally, we explored whether the flanking regions defined as 1 kbp up- and downstream of EWSR1-ETS-bound GGAA-mSats were enriched for transcription factor binding motifs (251 GGAA-mSats were bound by EWSR1-ETS in each cell line and examined for transcription factor motifs; 4,934 GGAA-mSats were not bound by EWSR1-ETS in any cell line and used as background in the analysis) by *de novo* motif finding with HOMER^66^. Across the ESCLA, the only enriched motifs matching known transcription factor binding motifs and fulfilling stringent selection criteria (1. not indicated as possibly false positive by the motif caller; 2. at least around 5% of targets as recommended in the motif caller’s description; 3. score for known motif at least default threshold 0.6; 4. minimum 5-fold enrichment) were those for *DUX4*, *GATA2/6* and *FOXF2* downstream, and *RUNX1* and *MIX1* upstream of the mSats (**Fig. 3d**). Of note, these transcription factor motifs were only present in about 5–7% of the tested target regions and solely *GATA2* appeared to be relatively highly expressed and EWSR1-ETS driven in the ESCLA, which may suggest that these transcription factors play a subordinate role compared to EWSR1-ETS in EwS pathogenesis.

We next utilized our ESCLA data to analyse the distance of EWSR1-ETS-bound and -unbound GGAA-mSats to EWSR1-ETS regulated genes across cell lines. As expected from the known enhancer-like activity of GGAA-mSats in EwS^19^, we noted a significantly (*P*<0.0001, Wilcoxon rank-sum test) shorter distance of the closest GGAA-mSat to the TSS of a given EWSR1-ETS induced gene, which was much longer in unregulated or suppressed genes (average distance 135 kbp versus 228 kbp and 461 kbp, respectively; **Fig. 3e**). This effect was even more prominent, when focussing only on EWSR1-ETS-bound GGAA-mSats (average distance 302 kbp versus 420 kbp and 415 kbp, respectively, *P*<0.0001; **Fig. 3f**). Vice versa, the TSS of EWSR1-ETS-repressed genes were significantly closer to EWSR1-ETS-bound sites located devoid of a GGAA-mSat (mostly single GGAA-motifs) than those of EWSR1-ETS-induced genes (average distance 363 kbp versus 402 kbp, *P*<0.0001). In line, EWSR1-ETS-repressed genes were enriched for nearby (2 Mbp around TSS) EWSR1-ETS-binding devoid of a GGAA-mSat compared to -induced genes (16.84 versus 13.65 binding events, *P*<0.0001), suggesting that repressed genes are likely controlled by EWSR1-ETS through binding to single ETS-like motifs.

To explore possible mechanisms of indirect EWSR1-ETS-mediated gene regulation, we tested the hypothesis that the promoters of such indirectly regulated genes should be enriched for binding motifs of EWSR1-ETS-regulated secondary transcription factors, using HOMER for 1 kb upstream to 50 bp downstream of the TSS of genes regulated in at least one third of cell lines upon EWSR1-ETS knockdown versus genes never found to be regulated by EWSR1-ETS (880 down-, 643 up-, and 15,967 unregulated genes). Although the promoters of the majority of EWSR1-ETS-regulated genes did not show evidence for enriched secondary transcription factor binding sites, we noted a significant enrichment of NFAT5 binding motifs in 12.8% of the EWSR1-ETS suppressed genes (versus 4.3% in unregulated genes). Conversely, in the promoters of EWSR1-ETS induced genes, an NFY or E2F1/2/4-like binding motif enrichment was observed in 27.7% and 11.1%, respectively (compared to 15% and 4.1% in unregulated genes, respectively) (**Fig. 3g**). Consistent with the hypothesis of a co-activating transcriptional function of these genes, *NFAT5* was downregulated by EWSR1-ETS in 3 of 18 EwS cell lines, whereas *NFYC* and *E2F1/2/4* were induced by EWSR1-ETS in 6, 12, 15 or 1 of 18 EwS cell lines, respectively. Furthermore, *E2F* gene regulation by EWSR1-ETS is in line with previous reports and our transcriptional core-signature comprising *E2F2* (**Suppl. Data 12**)^59, 67^.

### Association of heterogeneously EWSR1-ETS regulated genes with patients’ survival

As noted earlier, EwS is a clinically very heterogeneous disease^8^. Given the rather ‘silent’ genome and homogeneous epigenome of EwS^11–14, 56^ as well as the inherent polymorphic nature of GGAA-mSats, which can be fully recapitulated in our ESCLA, it is tempting to speculate that part of the observed clinical heterogeneity may be caused by the inter-individually variable composition of EWSR1-ETS-bound GGAA-mSats and thus differential EWSR1-ETS-mediated gene regulation. To investigate this possibility, we tested our list of heterogeneously EWSR1-ETS regulated genes (defined as being regulated strongly and weakly/not in at least one third of the 18 ESCLA cell lines, each; **Suppl. Data 14**) for association with overall survival in a cohort of *n*=196 EwS patients using the Mantel-Haenszel test. After Bonferroni-correction for multiple testing and filtering for genes being induced by EWSR1-ETS, we identified 19 heterogeneously regulated genes as being significantly associated with patients’ overall survival (**Table 1**). Strikingly, this approach highlighted the transcription factor *MYBL2* that was previously identified to be heterogeneously regulated by EWSR1-FLI1 due to germline variation in a proximal enhancer-like GGAA-mSat for which a higher number of consecutive GGAA repeats was positively correlated with MYBL2 expression in EwS cell lines and primary tumors^20^.

**Table 1.** Genes heterogeneously regulated in the ESCLA associated with overall survival. P-values are Bonferroni corrected; Surv. corr: expression status correlating with worse survival; IQR perc.: percentile of the inter-quartile range in the expression data of the survival cohort.

| Gene | Description | *P*-value | Surv.<br>corr. | IQR<br>perc. |
| --- | --- | --- | --- | --- |
| *AIF1* | Actin binding protein regulating immune cells | 0.0010 | high | 88 |
| *CDC20* | Cell cycle regulator | 0.0004 | high | 91 |
| *CDC25A* | Dual-specific phosphatase, DNA damage responder | 0.0028 | high | 50 |
| *CDC25C* | Cell cycle regulator, triggers mitosis | 0.0194 | high | 88 |
| *CENPI* | Centromere protein; E2F target | <0.0001 | high | 79 |
| *DUSP26* | Dual-specific phosphatase | 0.0161 | high | 66 |
| *E2F1* | Transcription factor, cell cycle regulator | 0.0001 | high | 34 |
| *EXO1* | Exonuclease | <0.0001 | high | 94 |
| *GPN3* | GTPase | 0.0002 | high | 68 |
| *GTSE1* | Cell cycle dependent protein | 0.0003 | high | 80 |
| *KIF14* | Microtubule regulator involved in mitosis | 0.0190 | high | 90 |
| *KIF15* | Microtubule regulator involved in mitosis | 0.0075 | high | 90 |
| *KIF2C* | Microtubule regulator involved in mitosis | <0.0001 | high | 88 |
| *MYBL2* | Transcription factor, involved in cell cycle | 0.0004 | high | 87 |
| *NEK2* | Protein kinase involved in mitosis | <0.0001 | high | 79 |
| *NUF2* | Centromere associated protein | 0.0011 | high | 97 |
| *SPAG5* | Associated with mitotic spindle apparatus | 0.0093 | high | 89 |
| *TPX2* | Microtubule regulator involved in mitosis | 0.0014 | high | 89 |
| *TRIP13* | ATPase involved in cell cycle progression | 0.0084 | high | 90 |

## DISCUSSION

Collectively, we generated the ESCLA – a hitherto unprecedented, comprehensive and multidimensional dataset comprising 18 well-curated EwS cell lines that were profiled at the genomic (WGS), epigenomic (ChIP-seq, methylation arrays), transcriptomic (DNA microarrays for gene expression and splicing), and proteomic level (mass spectrometry). Most of these high-throughput analyses were carried out in EE-high and -low conditions, which enabled the identification of previously unknown EWSR1-ETS regulated genes and proteins. Since all data were deposited at the NCBI Gene Expression Omnibus (GEO) or Sequencing Read Archive (SRA) and are freely available (accession codes PRJNA610192, GSE176339), we believe that this first version of the ESCLA will constitute an extremely rich resource, in particular for the EwS research community, and will also spur research directed at investigations of aberrant fusion transcription factor driven cancers in general. Furthermore, our targeted analyses of the ESCLA shed new light on the nature of EWSR1-ETS preferred GGAA-mSats, potential indirect modes of EWSR1-ETS-mediated gene regulation through secondary transcription factors, and highlighted putative co-regulatory transcription factor binding sites flanking EWSR1-ETS bound GGAA-mSats.

An important finding supported by our multidimensional ESCLA is that EwS cell lines are characterized by a duality of transcriptional stability on the one hand, which is mediated by largely EwS-specific genes, and plasticity on the other hand, which may be at least partially explained through the extraordinary variability of GGAA-mSats that serve as neo-enhancers for the EWSR1-ETS fusions. This duality may reason the so far experienced difficulties in identifying sensitive diagnostic marker-proteins for EwS^8^ and provided the basis for the identification of heterogeneous EWSR1-ETS target gene regulation of the current study, which may have implications for the clinical heterogeneity of EwS. These findings go beyond previous studies that have either analyzed single or just a few mostly EWSR1-FLI1-positive EwS cell lines by a limited set of technologies. Furthermore, the herein described duality underscores the importance of analyses based on multi-dimensional datasets of multiple cell lines, and may suggests that future functional studies in EwS could be more representative for EwS in general if being carried out in at least 3 different cell lines.

Among the limitations of the ESCLA, it should be noted that the read-length of the applied WGS procedure (150 bp paired-end) sets a barrier for calling the genotypes of especially long GGAA-mSats that may have thus escaped detectability and largely precluded robust conclusions on such potentially important EWSR1-ETS binding sites. Another important aspect to which our ESCLA falls short is the precise annotation of EWSR1-ETS regulated genes with the corresponding regulatory element that will be crucial to further dissect the regulatory landscape of EwS. Both aspects will constitute highly valuable extensions of the first version of the ESCLA that will be addressed using long-range sequencing technologies (e.g., Nanopore-sequencing) and chromosome conformation-capture methods (e.g., Hi-C). Furthermore, knockdown efficiency for EWSR1-ETS varied among the ESCLA cell lines, aggravating comparisons of DEGs and conflicting with a simple definition of DEGs with a uniform FC- and *P*-value-based threshold. To overcome these difficulties, we took the differential EWSR1-ETS knockdown efficiencies into account, defining DEGs with cell line-specific thresholds by an algorithm adapted from ROSE (see Methods and **Suppl. Fig. 2**). As a proof-of-concept, this cell line-specific approach identified and refined an EWSR1-ETS-dependent transcriptional core-signature (**Suppl. Data 12**), which contained a large number of previously identified EWSR1-ETS-regulated and functionally relevant genes^53, 55, 60, 62, 67^.

Nevertheless, our ESCLA constitutes – to the best of our knowledge – the largest and most comprehensive EwS cell line atlas that has been established to date. Thus, we believe that our study provides a blueprint for the systematic and multi-dimensional analysis of a large array of cancer cell lines of a specific cancer type, which can yield novel insights into the underlying principles of oncogene-mediated gene deregulation governing the complex and variable phenotype of a genetically homogenous but clinically heterogeneous disease.

## ABBREVIATIONS

ChIP-seq: chromatin immunoprecipitation with subsequent sequencing

DEG: differentially expressed gene

dox: doxycycline

EE-high/-low: EWSR1-ETS-high/-low

EEts: EWSR1-ETS

ESCLA: Ewing Sarcoma Cell Line Atlas

EF1: EWSR1-FLI1

EwS: Ewing sarcoma

FC: fold change

GSEA: gene set enrichment analysis

IHC: immunohistochemistry

IRS: immunoreactive score

mSat: microsatellite

SE: super-enhancer

shRNA: short hairpin RNA

TMA: tissue microarray

t-SNE: t-distributed stochastic neighbor embedding

TSS: transcription start site

WGCNA: weighted correlation network analysis

WGS: whole genome sequencing.

## AUTHORS’ CONTRIBUTIONS

MFO performed experimental assays. MFO, DS, and SZ performed ChIP. MFO performed bioinformatic analyses. AM and SO contributed to *in vitro* and *in vivo* experiments. FCA contributed to WCGNA analyses. SG did bioinformatic readout of ChIP-seq and SB the sequencing. JSG assembled the survival dataset and implemented the survival analysis tool. JA and AS provided additional EwS expression data with clinical annotation. KS, SMH, and ER performed MethylationEPIC BeadChip analyses and mass spectrometry. MS supported methylation data interpretation. SP supported WGS analyses. MG supported MFO in using HipSTR. TGPG, OD, and TK provided lab infrastructure. MFO and TG designed the display items and wrote the paper. TGPG conceived and supervised the study. All authors read the final version of this manuscript and approved it for submission.

## ACKNOWLEDGEMENTS

We thank Rebeca Alba Rubio, Stefanie Stein, Andrea Sendelhofert, and Anja Heier for excellent technical assistance, Mario Gipp and Heike Prelle for TMA construction, and Dr. Fabian Metzger for mass spectrometry readouts. We thank Dr. Paul Northcott and Dr. Brian Gudenas for providing R scripts for weighted correlation network analyses and Dr. Carlos Rodriguez-Martin for assistance with script adaptation. This project has been mainly supported by the German Cancer Aid (70112257 to TGPG). The laboratory of TGPG is further supported by grants from the LMU Munich’s Institutional Strategy LMUexcellent within the framework of the German Excellence Initiative, the ‘Mehr LEBEN für krebskranke Kinder – Bettina-Bräu-Stiftung’, the Wilhelm Sander-Foundation (2016.167.1), the Friedrich-Baur foundation, the Matthias-Lackas foundation, the Dr. Leopoldund Carmen Ellinger foundation, the Gert & Susanna Mayer foundation, the Deutsche Forschungsgemeinschaft (DFG 458891500), the German Cancer Aid (70114111, 70114278), the SMARCB1 association, the Boehringer-Ingelheim Foundation, and the Barbara & Wilfried Mohr Foundation. Further, this research was supported by the Helmholtz Zentrum München – German Research Center for Environmental Health, and within the Munich Center of Health Sciences (MC-Health), Ludwig Maximilian University, as part of LMUinnovativ. High-throughput sequencing of ChIP has been performed by the ICGex NGS platform of the Institut Curie supported by the grants ANR-10-EQPX-03 (Equipex) and ANR-10-INBS-09-08 (France Génomique Consortium) from the Agence Nationale de la Recherche (“Investissements d’Avenir” program), by the Canceropole Ile-de-France and by the SiRIC-Curie program – SiRIC Grant INCa-DGOS-4654.

## METHODS

### Provenience of cell lines

The EwS cell line A-673 (RRID CVCL_0080) was obtained from American Type Culture Collection (ATCC, Manassas, USA), and MHH-ES-1 (CVCL_1411), RD-ES (CVCL_2169), SK-ES-1 (CVCL_0627), SK-N-MC (CVCL_0530), TC-71 (CVCL_2213) were obtained from the German Collection of Microorganisms and Cell Cultures GmbH (DSMZ, Braunschweig, Germany). The EwS cell lines CHLA-10 (CVCL_6583), CHLA-25 (CVCL_M152), TC-32 (CVCL_7151) and TC-106 (CVCL_F531) were obtained from the Children’s Oncology Group (COG), and the EwS cell lines EW-1 (CVCL_1208), EW-3 (CVCL_1216), EW-7 (CVCL_1217), EW-22 (CVCL_1214), EW-24 (CVCL_1215), MIC (CVCL_EI96), Rh1 (CVCL_1658), and POE (CVCL_EJ01) were kindly provided by O. Delattre (Paris, France). HEK293T (CVCL_0063) was obtained from DSMZ. Cell identity was regularly controlled with in-house short tandem repeats (STR) profiling and, if applicable, by detection of specific fusion oncogenes by PCR, gel-electrophoresis and Sanger sequencing.

### Cell culture conditions

To avoid any bias due to culture conditions, all cell lines were cultured in the same medium, i.e. RPMI 1640 medium with stable glutamine (Biochrom, Berlin, Germany), supplemented with 10% FCS tested to be dox-free (Sigma-Aldrich, Taufkirchen, Germany) and penicillin and streptomycin (final concentrations 100 units/ml and 100µg/ml, respectively; Merck, Darmstadt, Germany) in tissue culture flasks and plates (TPP, Trasadingen, Switzerland). For improved attachment, for CHLA-10, CHLA-25, EW-3, EW-24, MIC, and TC-106 culture dishes were coated with 2% gelatine (Sigma-Aldrich) for cell expansion, and collagen type I solution from bovine skin (Sigma-Aldrich) in experimental assays. Cells were incubated at 37°C and 5% CO_2_ in a fully humidified environment. Cells were subcultured in ratios 1:2 to 1:8 after detachment with trypsin/EDTA (Biochrom), and spinning down the detached cells. Mycoplasma contamination was ruled out regularly by nested PCR with cell supernatant of each experiment.

### Establishment of stably transduced EwS cell lines with inducible knockdown of the fusion oncogene

18 EwS cell lines with inducible knockdown of the fusion oncogene were generated. The expression of an actual *EWSR1-ETS* gene fusion was assessed in all EwS cell lines by PCR of cell line cDNA and gel-electrophoresis. The following primers were used for *EWSR1*, *FLI1*, and *ERG*:

*EWSR1*-forward: 5’-GCCAAGCTCCAAGTCAATATAGC-3’;

*FLI1*-reverse: 5’-GAGGCCAGAATTCATGTTATTGC-3’;

*ERG*-reverse: 5’-TTGGGTTTGCTCTTCCGCTC-3’.

Commercial Sanger sequencing (MWG Eurofins genomics, Ebersberg, Germany) of the PCR products was performed to confirm the described transcript sequences at the fusion point of *EWSR1* and the *ETS* transcripts. For RNA interference (RNAi)-based knockdown of the fusion oncogene, the fusion-specific short hairpin RNA (shRNA) sequence published by Carrillo *et al.*^55^ was chosen for the 11 selected EwS cell lines with *EWSR1-FLI1* type 1 fusion (fusion of *EWSR1* exon 7 and *FLI1* exon 6). For four EwS cell lines with *EWSR1-FLI1* type 2 fusion (fusion of *EWSR1* exon 7 and *FLI1* exon 5), no 21 bases target sequence with at least 8 bases ranging from one fusion partner to the other yielded a high intrinsic score when tested in the Genetic Perturbation Platform (GPP, Broad Institute, Cambridge, USA). Most likely, this is caused by the last nucleotide of exon 7 of *EWSR1* being an adenosine, as is the last nucleotide of exon 4 of *FLI1*, and the first five nucleotides of exon 5 of *FLI1* being GTTCA as are the first nucleotides of exon 8 of *EWSR1*. Since these six nucleotides do not provide any specificity for RNAi, an shRNA predicted to be effective for *FLI1* exon 9 was chosen for knockdown experiments in these cell lines. Expression of wildtype (not-fused) *FLI1* was excluded for these EWSR1-FLI1 type 2 cell lines by qRT-PCR before application of this shRNA. The three *EWSR1-ERG* positive cell lines, CHLA-25, EW-3, and TC-106 exhibited all the same fusion transcript (*EWSR1* exon 7 fused to *ERG* exon 6). A fusion-specific shRNA target sequence for these *EWSR1-ERG* positive cell lines was predicted with GPP and specificity was tested with BLAST. Target sequences were as follows:

*EWSR1-FLI1* type 1: 5’-GCAGCAGAACCCTTCTTATGA-3’;

*EWSR1-FLI1* type 2: 5’-CTTTGGAGCCGCATCACAATA-3’;

*EWSR1-ERG*: 5’-GCTACGGGCAGCAGAATTTAC-3’;

Control shRNA: 5’-CAACAAGATGAAGAGCACCAA-3’.

All oligonucleotides were purchased from MWG Eurofins Genomics. For generation of cell liens with a dox-inducible knockdown of the respective *EWSR1-ETS* fusion oncogene, the lentiviral pLKO-Tet-On all-in-one vector^68^ (Addgene, Watertown, USA) was used. Via a tetracycline responsive element, the expression of the cloned shRNA can be induced by application of dox to the cell culture medium. This vector carries additionally a puromycin resistance cassette, enabling selection of successfully transfected cells. The vector was digested with EcoRI-HF and AgeI-HF (New England Biolabs, NEB, Frankfurt (Main), Germany), precipitated, and the opened plasmid without stuffer was extracted from agarose gel. The linearized vector was ligated with the annealed shRNA targeting the *EWSR1-ETS* fusions or with non-targeting control sequence using T4 DNA ligase (Thermo Fisher, Waltham, USA). Stellar competent cells (Clontech, TaKaRa, Saint-Germain-en-Laye, France) were transformed with the ligated vector and plated. Bacteria clones were picked, and presence of the vector was controlled with colony PCR. Corresponding clones were expanded, plasmids were extracted (PureYield Plasmid Midiprep System, Promega, Maddison, USA) and correct sequence of the shRNA was controlled with sanger sequencing (MWG Eurofins Genomics). Lentiviral particles were generated by transfecting HEK293T cells with 10 µg shRNA-carrying plasmid, 10 µg D8.9 and 3 µg pCMV-VSV-G (Addgene) packaging plasmids using Lipofectamine LTX with Plus Reagent (Thermo Fisher). The standard culture medium was replaced after 12 h by culture medium with 30% FCS. Virus-containing supernatant was collected after 48 h and filtered through 0.45 µm membrane. About 1×10^6^ EwS cells were infected with 1 ml supernatant without polybrene. Upon confluence of the cells, cells were split, and successfully stably transduced cells were selected with 1 µg/ml puromycin (InvivoGen, San Diego, USA). This concentration has been tested to be the lowest lethal dose for these EwS cell lines in wildtype conditions. For single cell cloning, virtually 0.8 cells per well of a 96-well-plate were seeded in 150µl medium, containing 20% conditioned medium of the respective cell lines filtered through 0.45 µm membrane. The cloned shRNAs against the respective *EWRS1-ETS* fusion oncogenes was induced by addition of 1 µg/ml dox (Merck, Darmstadt, Germany) to the cell culture medium. Dox was refreshed every 48 h.

### Expansion of EwS cell lines *in vivo*

Animal experiments were conducted with allowance from the government of Upper Bavaria (ROB-2532.Vet_02-15-184) in the proprietary animal facility of the Institute of Pathology, LMU Munich, and in compliance with all relevant ethical regulations. Sample size was predetermined using power calculations with *β*=0.8 and *α*=0.05 based on preliminary data and in compliance with the 3R system (replacement, reduction, refinement). To generate EwS xenograft tumors for immunohistochemistry, 2.5×10^6^ cells of six representative EwS cell lines with inducible knockdown of the fusion oncogene (A-673, EW-7, POE, SK-N-MC, Rh1, TC-71) were injected in 1:1 mix of PBS (Biochrom) and basement membrane matrix (Thermo Fisher) into the right flank of NOD/Scid/gamma mice (NSG, Charles River, Wilmington, USA). Mice were controlled every two days for health and tumor growth. Tumor size was measured with a caliper and tumor volume calculated as *V=a×b²/2*, with *a* being the largest and *b* the smallest diameter. Once, the average tumor diameter reached 10 mm, mice were treated for 96 h with 2 mg/ml dox (belapharm, Vechta, Germany) in sucrose (Sigma-Aldrich) drinking water or only sucrose drinking water, and then sacrificed by cervical dislocation. Other humane endpoints were determined as follows: Xenograft volume of 1,500mm³, ulcerated tumors, loss of 20% body weight, constant curved or crouched body posture, bloody diarrhoea or rectal prolapse, abnormal breathing, severe dehydration, visible abdominal distention, obese Body Condition Scores (BCS), apathy, and self-isolation. Xenografts were quickly extracted. Fragments of the tumors were snap frozen for RNA isolation. The remaining tumor was fixed in formalin and, at latest 72 h later, dehydrated and embedded in paraffin.

### RNA extraction, reverse transcription and quantitative real-time polymerase chain reaction (qRT-PCR)

RNA was extracted from cells with the NucleoSpin RNA kit from Macherey-Nagel (Düren, Germany) following the manufacturer’s protocol. RNA (1 µg, quantified with NanoDrop) was reverse transcribed to cDNA with the High-Capacity cDNA Reverse Transcription Kit (Thermo Fisher) according to the manufacturer’s protocol. cDNA was diluted 1:10. For qRT-PCR 6.75µl cDNA with 7.5 µl SYBR Select Mastermix (Thermo Fisher) and 0.75 µl equimolar mix of forward and reverse primer (final concentration 0.5 µM) were mixed per reaction (final volume 15 µl). Each reaction was run in duplicates. Gene expression levels for the tested gene were normalized to the expression of the housekeeping gene *RPLP0*, which has been previously used for EwS cells^18, 20, 41^ and that does not show significant variation of its expression levels upon knockdown of the respective fusion gene as determined by Affymetrix Clariom D microarrays. qRT-PCR was run in a CFX Connect Real-Time PCR Detection System (Bio-Rad, Munich, Germany) with the following protocol: heat activation at 95°C for 2 min, annealing and elongation at 60°C for 30 sec (50 cycles), final denaturation at 95°C for 30 sec, followed by stepwise temperature increase from 65°C to 95°C, 0.5°C step every 5 sec, for melting curve. Differential expression in the qRT-PCR samples was assessed with the Delta-Delta-Ct method. Primer sequences were as follows:

*RPLP0* forward: 5’-GAAACTCTGCATTCTCGCTTC-3’;

*RPLP0* reverse: 5’-GGTGTAATCCGTCTCCACAG-3’;

*EWSR1-FLI1* forward: 5’-GCCAAGCTCCAAGTCAATATAGC-3’;

*EWSR1-FLI1* reverse: 5’-GAGGCCAGAATTCATGTTATTGC-3’;

*EWSR1-ERG* forward: 5’-TCCAAGTCAATATAGCCAACAGAG-3’;

*EWSR1-ERG* reverse: 5’-CTGTGGAAGGAGATGGTTGAG-3’;

*EWSR1* (wildtype) forward: 5’-CAGCCAAGCTCCAAGTCAATA-3’;

*EWSR1* (wildtype) forward: 5’-TCCAGACTCCTGCCCATAAA-3’;

*FLI1* (wildtype) forward: 5’-TGGATGGCAAGGAACTGTG-3’;

*FLI1* (wildtype) reverse: 5’-CGGTGTGGGAGGTTGTATTA-3’;

*ERG* (wildtype) forward: 5’-CGAACGAGCGCAGAGTTAT-3’;

*ERG* (wildtype) forward: 5’-ACGTCTGGAAGGCCATATTC-3’. Primers for the reverse fusion oncogene were published before ^26^.

### Western blot

About 1×10^6^ EwS cells were lysed in 100 µl RIPA buffer. Protein content was measured with Bradford assay (Bio-Rad). 20 µg protein were denaturated at 95°C for 5 min, and loaded on western blot gel. Samples were separated in a 10% acrylamide (Roth, Karlsruhe, Germany) gel and blotted onto nitrocellulose membrane (GE Healthcare, Freiburg, Germany) in a wet system. The membrane was blocked with milk (Roth) or BSA (for anti-FLI1) and incubated with the first antibody overnight at 4°C. Incubation with the corresponding horseradish peroxidase coupled second antibody followed the next day for 1 h at room temperature. Bands were detected and quantified after addition of a chemiluminescent reagent (Merck) with LI-COR Odyssey (Homburg, Germany). Signal intensities were automatically quantified by the LI-COR system. The following antibodies were used: monoclonal anti-FLI1 raised in mouse (254M-16, 1:1,000; medac, Wedel, Germany), monoclonal anti-ERG raised in rabbit (EP111, 1:2,000; Cell Marque, Rocklin, CA, USA), anti-GAPDH raised in mouse (sc-32233, 1:2,000; Santa Cruz, Dallas, USA), polyclonal anti-mouse IgG-HRP raised in goat (W402B, 1:3,000; Promega) and polyclonal anti-rabbit IgG-HRP raised in goat (R1364HRP, 1:5,000; OriGene, Herford, Germany)

### Tissue microarray (TMA) construction and immunohistochemistry (IHC)

On hematoxylin & eosin (HE) stained slides of the EwS xenograft tumors grown in NSG mice, representative areas with vital tumor tissue were marked. Three tissue cores (each 1 mm in diameter) per tumor were extracted from these areas. Tissue cores were integrated in TMA scaffolds with human tonsils as control tissue. For subsequent IHC stains, 4 µm sections were cut. Antigen retrieval was performed with microwave treatment using the antigen retrieval AR-10 solution (HK057-5K, DCS Innovative, Hamburg, Germany) for FLI1, the antigen retrieval ProTaqs I and V Antigen-Enhancer (Quartett, Berlin, Germany) for p-MYBL2 and PAX7, and the Target Retrieval Solution (S1699, Agilent Technologies, Waldbronn, Germany) for SOX6. For blockage of endogenous peroxidase, slides were incubated for 20 min in 7.5% aqueous H_2_O_2_ solution and blocking serum. Then, slides were incubated for 60 min with the primary anti-FLI1 (254M, 1:120; Cell Marque, Rocklin, USA), anti-p-MYBL2 (ab76009, 1:100; Abcam, Cambridge, UK), anti-PAX7 (PAX7-c, 1:180; DSHB, Iowa City, IA, USA) or anti-SOX6 (HPA003908, 1:1,600; Atlas Antibodies, Stockholm, Sweden) and afterwards with a secondary peroxidase-conjugated anti-rabbit/-mouse IgG antibody (MP-7401, ImmPress Reagent Kit for FLI1, p-MYBL2 and SOX6, PK6200, Vectastain Elite ABC HRP Kit for PAX7; Vector Laboratories, Burlingame, USA). Target detection was performed using DAB+chromogen (Agilent Technologies, Santa Clara, USA). Slides were counterstained with hematoxylin Gill’s Formula (H-3401; Vector Laboratories). Immunoreactivity was semi-quantified with a slightly modified immunoreactivity score by Remmele and Stegner^69^. Staining intensity was scored as no, low, moderate, and strong with the values 0, 1, 2, 3, the area of stained cells was scored with 0 to 5 indicating quintiles of cell area positive for immunoreactivity. The product of the intensity and the area score resembled the final immunoreactivity score (IRS).

### Whole Genome Sequencing (WGS)

DNA was extracted from wildtype EwS cell lines using the NucleoSpin Tissue kit (Macherey Nagel) following manufacturer’s protocol and eluted in H_2_O. For sequencing, 50 µl of 50 ng/µl DNA were used. After initial DNA quality assessment on a bioanalyzer (DNA Integrity Number at least 7.0), DNA was sequenced on Illumina HiSeq Xten (150 bp, paired-end; Illumina, San Diego, USA) and a PCR-free protocol at the Genomics and Proteomics Core Facility of the German Cancer Research Center (DKFZ, Heidelberg, Germany). Raw sequencing data were aligned to the human reference genome (version hg19) with Burrows-Wheeler Aligner (bwa mem)^70^. PhiX contamination was excluded, Illumina adapters and duplicates were marked with picard (Broad Institute). Base quality scores were recalibrated with GATK (Broad Institute). Quality of the final alignments was controlled with FASTQC^71^. Coverage was assessed with samtools depth^72^ and displayed as average of 90 kb bins. *De novo* motif finding was performed with HOMER^66^.

### Genotyping of GGAA-mSats

To genotype and phase GGAA-mSats in EwS cell lines, the haplotype inference and phasing for short-tandem repeats (HipSTR) tool was employed^30^. A library of all potential GGAA-mSats was generated with Tandem Repeats Finder^65^ running on the human reference genome (hg19), calling for those repeats with four nucleotide motifs, either high guanine-adenine or cytosine-thymine content and at least four sequential GGAA or TTCC motifs. The final library contained 8,311 loci. The library, the processed and aligned Illumina WGS data, and the reference genome were used as input for HipSTR. HipSTR mines all STR alleles for each locus, aligns reads accounting for artefacts due to the diversity and structure of STRs, and phases the STRs. The resulting variant calling file (VCF) was filtered for minimal call quality (0.9), maximum number of stutter artefacts and indels in STR flanking regions (0.15), minimal call rate (0.3), minimal call depth per locus (10) and minimal supporting reads per allele (3). Readouts were displayed for up to 18 consecutive GGAA-repeats as up to this repeat number more than 100 genotypes were called.

### Copy number analysis

Copy numbers of genomic regions were inferred from the WGS data with CNVnator^73^ by extraction of read mapping information, building a read depth histogram, calculating statistics and CNV calling with a bin size of 1,000 nt. CNVkit was employed for CNV analyses for single genes with 300 bp bind and exclusion of not accessible chromosomal regions^74^.

### Variant calling

Single nucleotide variants (SNVs) were called with bcftools^75^ and GATK Mutect2 for somatic mutations, and annotated with ANNOVAR and SnpEff^76, 77^. Structural variants were assessed with LUMPY, giving split and discordant reads as input, and GRIDSS^78–80^. Transchromosomal fusions were analysed with BreakDancer^27^. Genomic rearrangement loops were extrapolated using the CNVkit and GRIDSS output in ChainFinder^25^.

### Chromatin immunoprecipitation with subsequent next generation sequencing (ChIP-seq)

To identify EWSR1-ETS-bound genomic regions, ChIP was performed. For ChIP, at least 1×10^7^ EwS cells were cultured. Only cells from not confluent cultures without consumed culture medium were used. Cells were fixed with 1% methanol-free formalin for 10min at RT. Formalin was quenched with glycine (final 200 mM) for 5 min. Cells were lysed in two steps with the Diagenode iDeal ChIP-seq kit for transcription factors (Diagenode, Seraing, Belgium).

DNA was sheared with Bioruptor Plus sonication device (Diagenode). Cell debris was spun down, and sheared DNA was added to antibodies coupled to washed magnetic beads (anti-FLI1 2 µg, ab15289, Abcam; anti-ERG 2 µg, ab92513, Abcam; anti-H3K4me3 1.4 µg, C15410003, Diagenode; H3K27me3 2.9 µg, C15410069, Diagenode; H3K27ac 1µg, ab4729, Abcam). As wildtype FLI1 or ERG are not expressed in EWSR1-FLI1 or EWSR1-ERG positive EwS cell lines, respectively, anti-FLI1 and anti-ERG antibodies can be assumed to specifically target the fusion oncoprotein. After incubation overnight at 4°C, beads were washed 4× with Diagenode washing buffers, bound DNA was eluted from the beads and purified. DNA was quantified with Qubit. Ten µg DNA were sequenced on Illumina HiSeq2500 platform (100 bp or 150 bp, single-end). As control for successful immunoprecipitation and suitability of the selected antibodies ChIP-PCR was performed with primers for genomic regions known to be bound by the respective marker. Raw sequencing data were aligned to human reference genome hg19 with bowtie2^81^. Alignments were purged from duplicates with samtools. Peaks were called with MACS2^82^, selecting narrow peak for transcription factors and broad peak for histone marks. Super-enhancers were called with ROSE^57^.

For data display in IGV^83^, MACS2 output was converted into bigwig files. These files were normalized between cell lines based on genes with steady expression levels across cell lines using CHIPIN (https://github.com/BoevaLab/CHIPIN). Merged peaks of ETS-ChIP across cell lines were calculated with HOMER and 200 nt width around peak center. For merged H3K27ac peaks, peak files were used as given by MACS2. Overlaps and genomic distances were analyzed with BEDTools^84^. Of note, for distance analyses to EEts-binding sites.

### DNA microarray expression gene analyses

For identification of EWSR1-ETS regulated genes, DNA microarray gene expression analyses were performed. From all 18 EwS cell lines with inducible knockdown of the respective fusion oncogene, cells were seeded in three technical replicates in standard medium or standard medium supplemented with dox (1 µg/ml). Medium was changed to fresh medium (including dox) after 48 h. After 96 h, cells were lysed and RNA was extracted with the NucleoSpin RNA kit (Macherey Nagel). Knockdown of the respective fusion oncogene was evaluated by qRT-PCR. Quality of the RNA (1 µg) was controlled (RNA Integrity Number ≥ 9.0 in all samples) before reverse transcription, fragmentation and hybridization with the Affymetrix Clariom D microarray (Thermo Fisher) at IMGM (Planegg, Germany). The resulting raw data (CEL files) were normalized and summarized with the SST-RMA analysis algorithm^85^ in the Transcriptome Analysis Console (version 4.0, Thermo Fisher) and manufacturer’s array description file (version 2). Gene expression values were log2-transformed. Readouts not corresponding to actual gene transcripts were filtered out.

Differentially expressed genes (DEGs) were defined like super-enhancers by the ROSE algorithm instead of application of arbitrary fold change and *P* value cut-offs. Specifically, genes were plotted ranked by their mean expression fold change in wildtype versus knockdown condition. Linear functions connecting the lowest fold change with the value 0, and connecting the highest fold change with the value 0 were calculated. The *y*-intercept for the linear functions were adapted until the function for the positive and negative fold changes had only one intersection point with the plotted positive and negative fold changes, respectively. All genes from the intersection point to the extremes were defined as DEGs (**Suppl. Fig. 2**). Those genes were depicted as heterogeneously regulated, which were in at least 6 cell lines strongly regulated (top 33% of regulated genes) and in at least another 6 cell lines weakly (lower 33% of regulated genes) or not regulated. Overlaps of regulated genes were calculated for each cell line versus all others. Only the top 924 regulated genes (minimal number of genes defined as regulated in a single cell lines) were considered. To increase overlap rates, the top 33% of the 924 regulated genes per cell line were compared to the top 924 regulated genes of each other cell line. Nonlinear dimensionality reduction was used to examine potential EwS cell line clusters dependent on the respective fusion oncogene. To this end, gene expression values on the transcriptome level in EWSR1-ETS-high condition were analyzed with the Rtsne package (version 0.15; perplexity=5; max_iter=500).

### Protein quantification

For identification of EWSR1-ETS regulated proteins, mass spectrometry on protein lysates of EwS cell lines was performed. From all 18 EwS cell lines with inducible knockdown of the fusion oncogene, cells were seeded in three technical replicates in standard medium or standard medium supplemented with dox (1 µg/ml). Medium was changed to fresh medium after 48 h (including dox). After 96 h, the cell surface was washed with FCS-free medium, before incubation in FCS-free medium for 20 min and a second wash step to clean the cells from serum proteins. Cells were lysed in Nonidet-P40 buffer (1 % Nonidet P40 (Thermo Fisher) and complete protease inhibitor (Sigma-Aldrich)). Lysates were collected with cells scrapers into protein low-binding tubes and sonicated with 60 % amplitude, 6× 30 s. Protein content of the lysates was quantified with Bradford assay (Bio-Rad) and 10 µg proteins were proteolysed with trypsin and peptides analysed by quantitative LC-MSMS on a Q Exactive HF (Thermo Fisher) by a data-independent acquisition approach as described earlier^86^. Expression of 3,248 proteins were quantified consistently across all 18 cell lines. The expression levels of additional 1,336 proteins were imputed from patchy data with a machine learning based algorithm implemented in fancyimpute (v. 0.5.4, IterativeSVD option). The Pearson’s correlation coefficients in the smaller dataset versus the imputed one were roughly the same with 0.56 versus 0.58, respectively, both highly significant. Proteins regulated upon EWSR1-ETS knockdown were defined as genes on the transcriptome level. Overlaps of regulated proteins were calculated for each cell line versus all others. Only the top 216 regulated proteins (minimal number in a single cell lines) were considered. To increase overlap rates, the top 33% of the 216 regulated proteins per cell line were compared to the top 216 regulated proteins of each other cell line. Clustering of EwS cell lines, dependent on the respective fusion oncogene, was investigated using t-SNE analysis (see DNA microarray expression gene analyses).

### Methylation analysis

For analysis of EWSR1-ETS dependent CpG island methylation, genomic DNA of all 18 EwS cell lines with and without knockdown of the respective fusion oncogene was genotyped on Illumina Infinium MethylationEPIC BeadChip arrays. Therefore, all EwS cell lines were seeded in three technical replicates in standard medium or standard medium supplemented with dox (1µg/ml). Medium was changed to fresh medium after 48 h (including dox). After 96 h, samples were lysed and DNA was extracted with the NucleoSpin Tissue kit (Macherey Nagel). Genomic DNA was genotyped with the Illumina Infinium Methylation EPIC BeadChip arrays at the Molecular Epidemiology unit of the German Research Center for Environmental Health (Helmholtz Center, Munich, Germany). Readout and analysis of the EPIC arrays was performed with GenomeStudio (Illumina) and the R minfi package and bumphunter algorithm^87^. For t-SNE analysis, the same approach and tools as for transcriptome and proteome data were applied. Methylation profile classification was run on the MolecularNeuropathology.org website^56, 88^.

### Gene set enrichment analysis (GSEA), weighted correlation network analysis (WGCNA), and gene ontology (GO) analyses

To identify gene sets that are enriched among EWSR1-ETS regulated genes, all genes from DNA microarray expression analysis were ranked by their expression fold change in wildtype (fusion oncogene high) versus knockdown (fusion oncogene low) condition. Pre-ranked GSEA was performed with 1,000 permutations^89^ on previously described gene sets obtained from the Molecular Signature Database hosted at the Broad Institute (MSigDB, c2.all.v6.2). For network analysis, the weighted correlation network analysis R package (WGCNA R)^90^ was employed. A matrix of functionally annotated gene sets versus genes (indicating presence of the gene in the respective set) was built and the Jaccard’s distance for all possible pairs was computed generating a symmetric GSEA adjacent matrix. The dynamicTreeCut algorithm was employed to identify GSEA term clusters. Top results were selected for visualization (min absolute NES=2.5). The cluster label corresponds to the highest scoring node of each cluster. For network visualization Cytoscape (v 3.8.2) was used^91^. Selected gene lists were probed for enriched gene ontology terms using PANTHER^92^.

### Survival analysis

For identification of genes associated with overall survival of EwS patients, an established dataset^93^ composed of gene expression microarray data of 196 primary EwS tumors with clinical annotation available at GEO (accession codes: Affymetrix HG-U133plus2: GSE12102, GSE17618, GSE34620; Affymetrix HuEX-1.0st: GSE63157) or obtained from J. Alonso (Amersham/ GE Healthcare CodeLink microarrays, unpublished data) was used. Signal raw data were normalized and summarized with Robust Multi-array Average (RMA)^85^ and custom brainarray chip description files (CDF, v20), yielding one optimized probe-set per gene^94^. Batch effects between microarray types were removed with ComBat^95^. Only samples with tumor purity of at least 60%, calculated with the ESTIMATE algorithm^96^, were further analyzed (TCGA standard). Survival association of the genes represented on all microarrays was calculated with the Mantel-Haenszel test using an in-house tool (GenEx) and GraphPad PRISM (version 8; GraphPad Software Inc., CA, USA). *P* values were adjusted for multiple testing using Bonferroni correction.

### Statistical tests and data visualization

Association of gene expression levels with patients’ overall survival was assessed using the Kaplan-Meier method and the Mantel-Haenszel test. For comparison of two groups with normal data distribution (as assessed by Kolmogorow-Smirnow test), the two-sided independent Student’s *t*-test was used. For statistical comparison of two groups in which normal distribution could not be assumed, the two-sided Wilcoxon rank-sum test was applied. For assessment of statistical significance assessment between two groups with two discrete categories, the Fisher’s exact test or chi-square test was used.

Readouts of ‘omics’ analyses for specific genomic regions were visualized in the Integrative Genomics Viewer (IGV; version 2.6.2). Data encompassing the entire human genome were displayed in Circos plots^97^. Heatmaps were generated with GEN-E (Broad Institute) and Venn diagrams in BioVenn. t-stochastic neighbor embedding was performed in R. Other plots were generated in GraphPad PRISM (version 8; GraphPad Software Inc) and R.

## Data and code availability

Original WGS, ChIP-seq, MethylationEpic BeadChip and DNA microarray data that support the findings of this study were deposited at the National Center for Biotechnology Information (NCBI) SRA and GEO, and are accessible through the bioproject and series accession numbers PRJNA610192 and GSE176339. All other data supporting the findings of this study are available within the article, its supplementary data files, or from the corresponding author upon reasonable request. Custom code is available from the corresponding author upon reasonable request.

## LEGENDS FOR SUPPLEMENTARY FIGURES

**Supplementary Figure 1.**
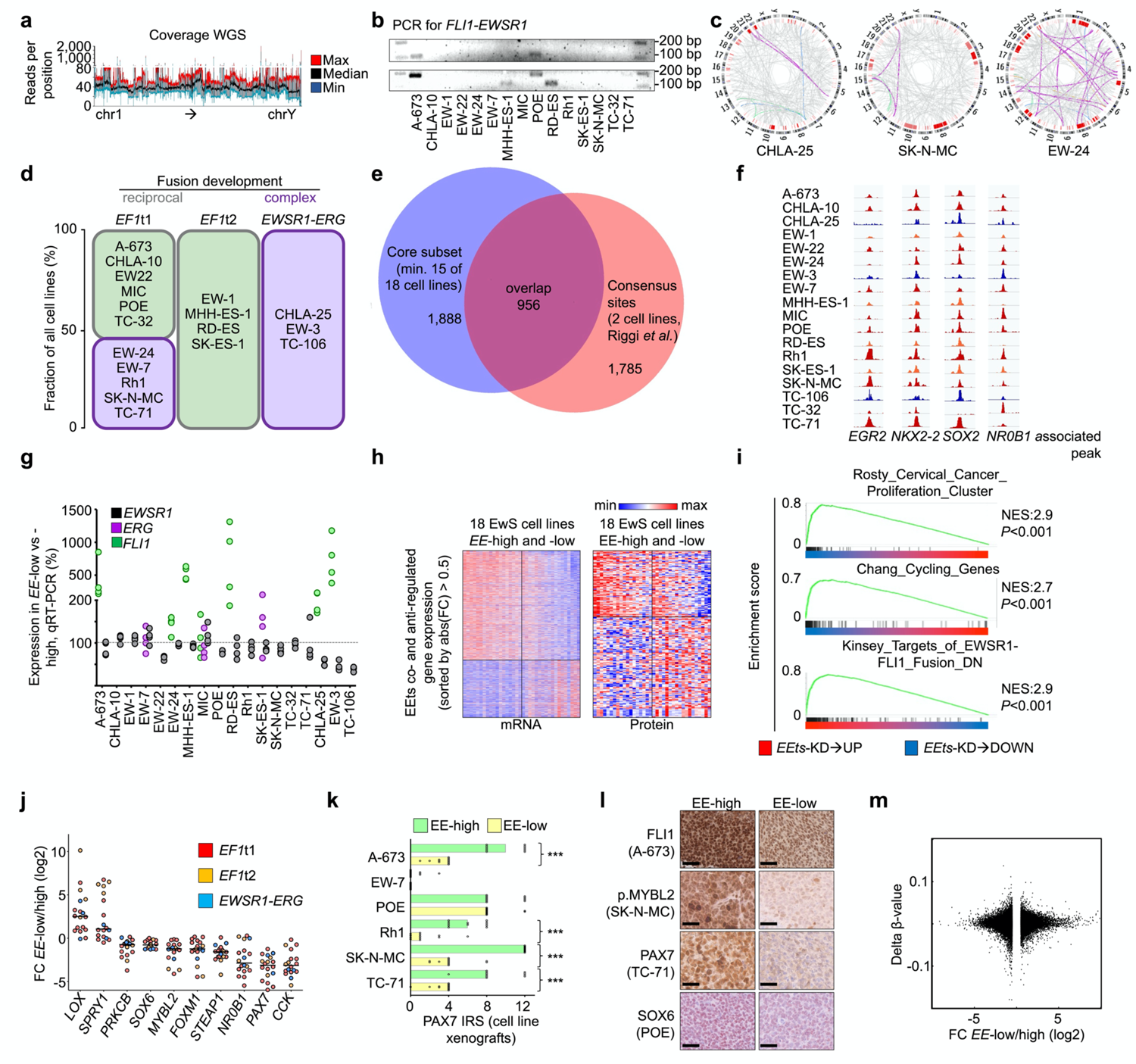

**a)** Coverage of ESCLA whole genome sequencing (WGS) data over the entire human genome counted per 90 kb window, maximum (max), median and minimum (min) coverage across cell lines, range is depicted in grey. **b)** Gel electrophoresis of the PCR amplified reverse *FLI-EWSR1* transcript as indicator for reciprocal translocation as mechanism of fusion genesis; 100 bp ladder. **c)** Circos plots indicating most significant genomic rearrangements per cell line in purple, including the *EWSR1-ETS* fusion. **d)** Overview of the ESCLA cell lines and the respective type of fusion development; *EF1*t1/2: *EF1* subtype 1 or 2. **e)** Venn diagram indicating the overlap of EWSR1-FLI1 consensus binding sites described by Riggi *et al.*^19^ and the core subset (EWSR1-ETS-bound sites in at least 15 of 18 EwS cell lines) from the ESCLA. **f)** IGV plots of EWSR1-ETS-ChIP-seq at known EWSR1-ETS targets regulating GGAA-mSats across all cell lines of the ESCLA. **g)** Dot-bar plot indicating differential expression of non-fused *EWSR1*, *ERG*, and *FLI1* in ESCLA cell lines in *EWSR1-ETS*-low versus -high (*EE*-low/high) condition (*n*=4, mRNA level, assessed by qRT-PCR). Only genes with expression of at least 1‰ of the fusion in *EE*-high state were considered as expressed and depicted in the figure. **h)** Heatmaps indicating strongly differentially expressed genes in EE-high and -low state of 18 EwS cell lines on mRNA and protein level. Rows represent genes with average absolute expression FC across cell lines higher than 0.5, sorted from top to bottom from lowest to highest average FC; columns represent the ESCLA cell lines in fusion oncogene-high (left) and -low (right) condition, sorted from left to right by descending average gene expression levels; EEts: EWSR1-ETS. **i)** Exemplary gene set enrichment analysis (GSEA) graphs for *EEts* co- and anti-regulated genes. NES: normalized enrichment score. **j)** Dot plot indicating the fold changes (FC) of the depicted EWSR1-ETS target gene’s expression on mRNA level upon EWSR1-ETS knockdown. **k)** Dot bar plot indicating PAX7 immunoreactivity as immunoreactive score (IRS) in EE-high and -low EwS cell line xenografts; one to three stars indicate significance level of 0.05, 0.01 and 0.001, respectively. **l)** Exemplary micrographs of immunohistochemically FLI1, p.MYBL2, PAX7 and SOX6 stained EwS cell line xenografts in EE-high and -low condition; scale bar indicates 50 µm. **m**) Scatter plot indicating genes with absolute FC on the RNA-level in *EE*-low vs. -high condition >0.5 in more than one third (>6) of the ESCLA cell lines and the average delta of the β-values of CpG sites in *EE*-low vs. -high condition, which were uniquely annotated for the promoters of the respective gene.

**Supplementary Figure 2.**
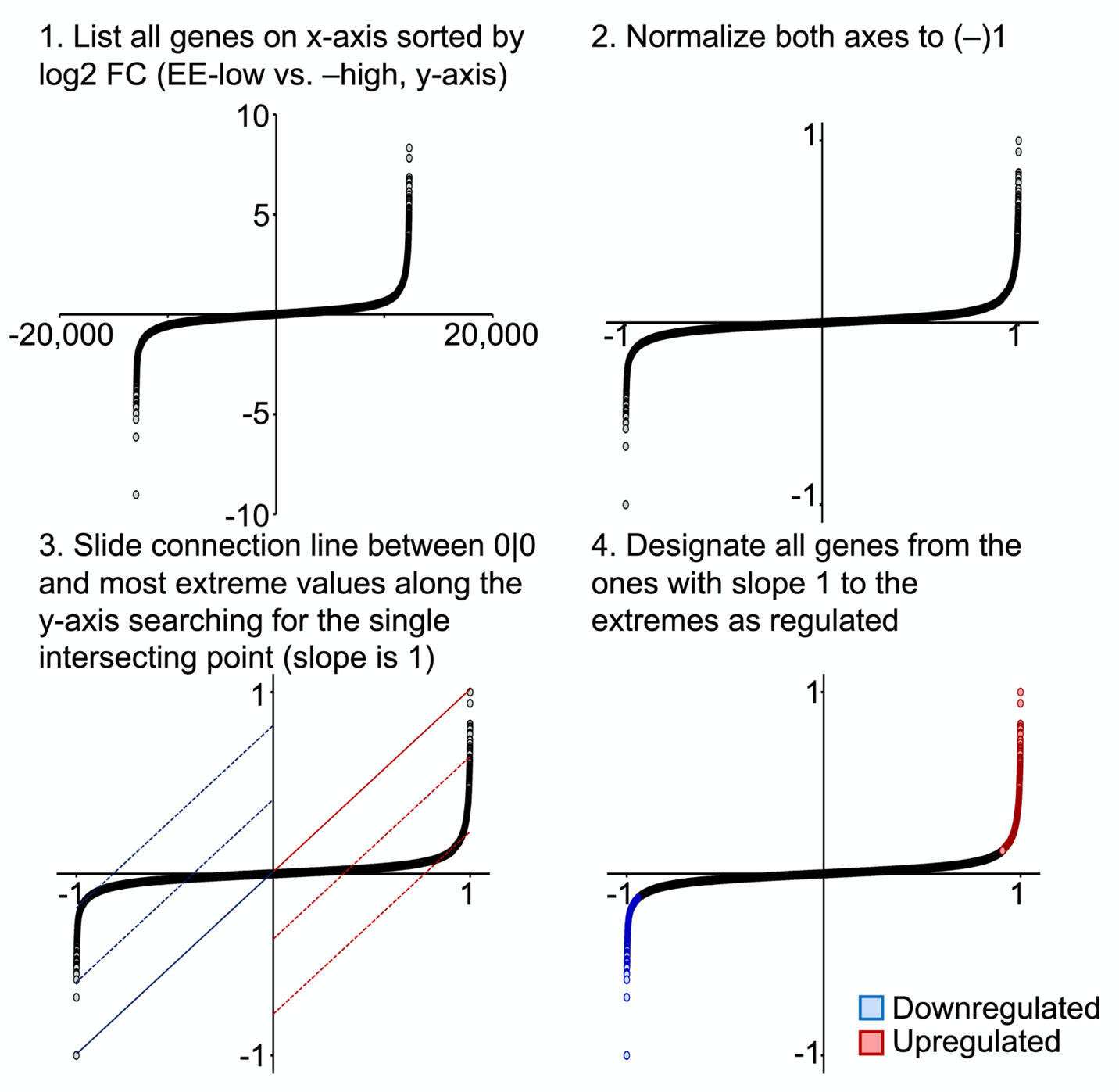
Workflow for the designation of regulated genes applied per cell line on the transcriptome and proteome level of the ESCLA to avoid arbitrary cut-offs and *P*-value based approaches, and account for differential EWSR1-ETS knockdown efficiencies. EE-low/-high: EWSR1-ETS-low/-high; FC: fold change.

